## Supplementary Figures for "LINE-1 replication in a mouse TDP-43 model of neurodegeneration marks motor cortex neurons for cell-intrinsic and non-cell autonomous programmed cell death"

Supplementary Fig 1a

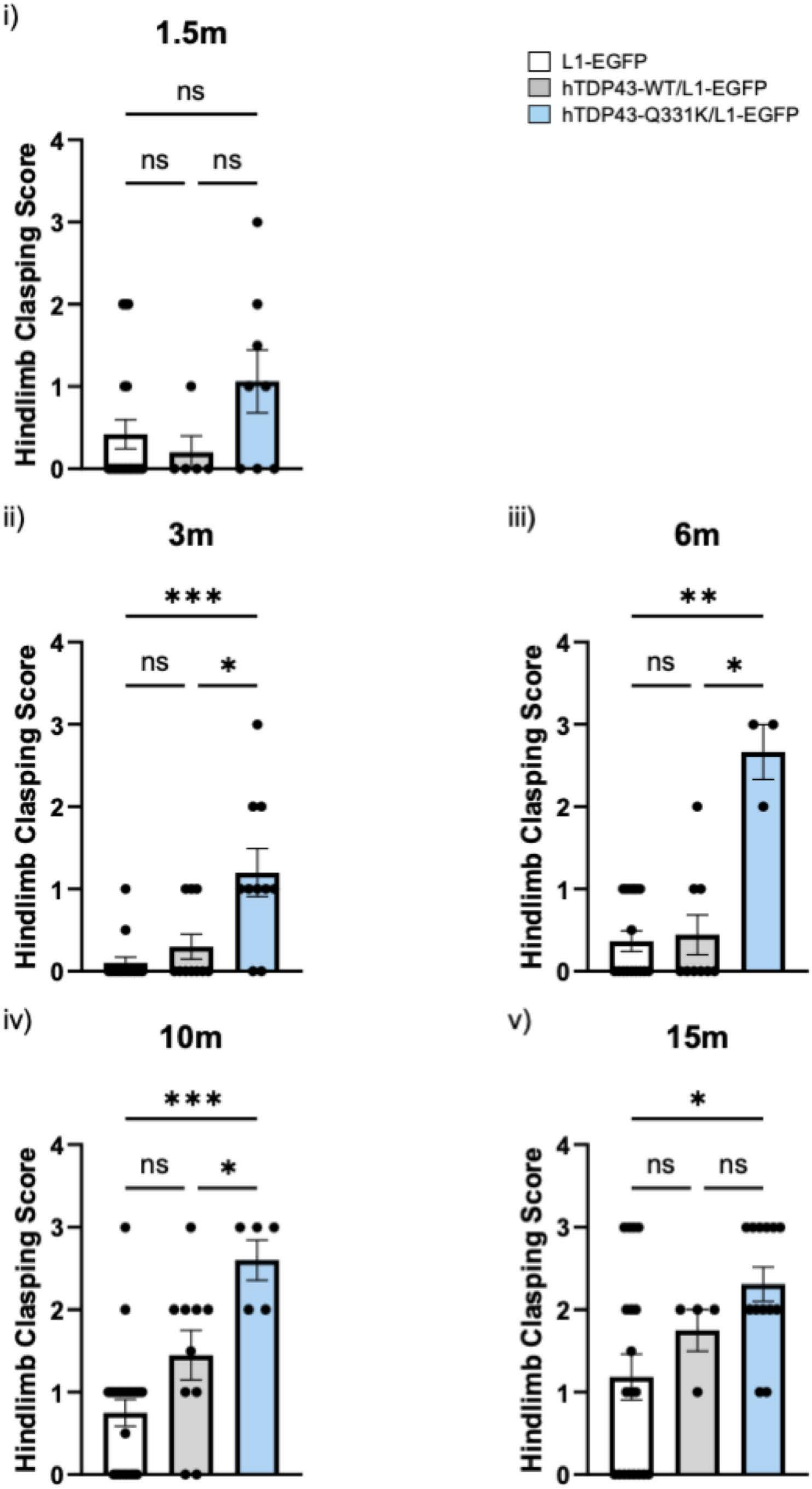

#### i) Differential expression of all RTE

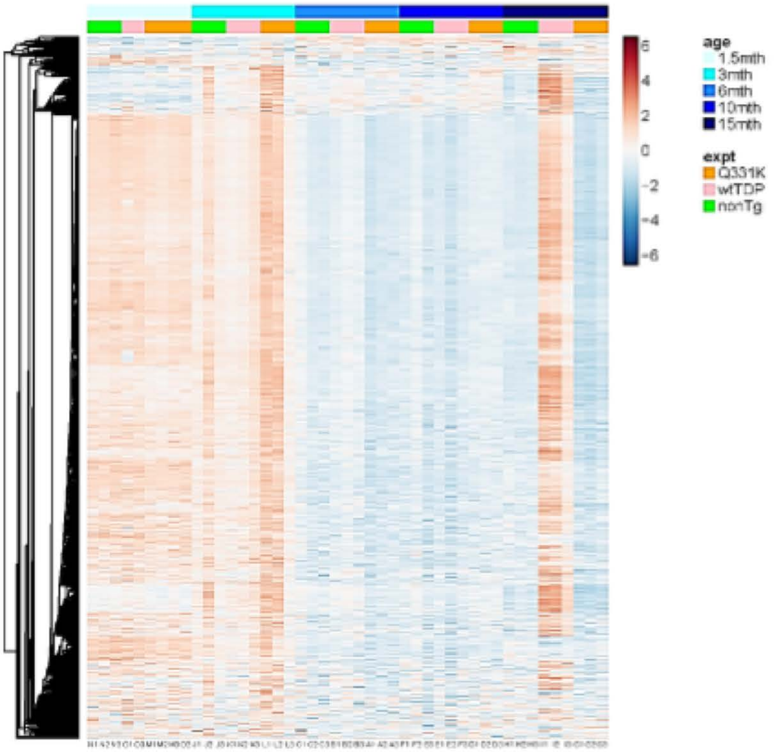


#### ii) Significantly expressed genes and RTE using GLM

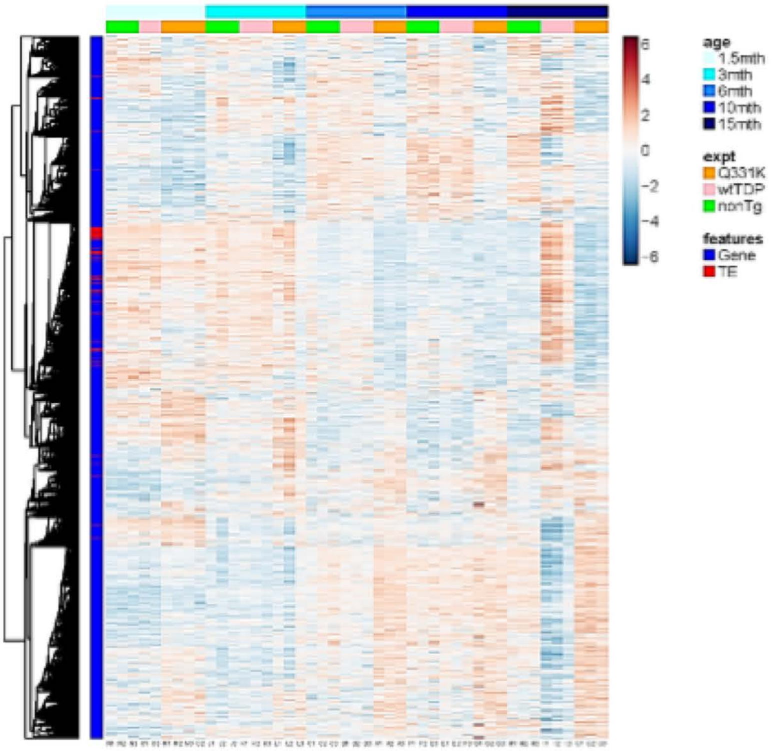

Supplementary Figure 2b

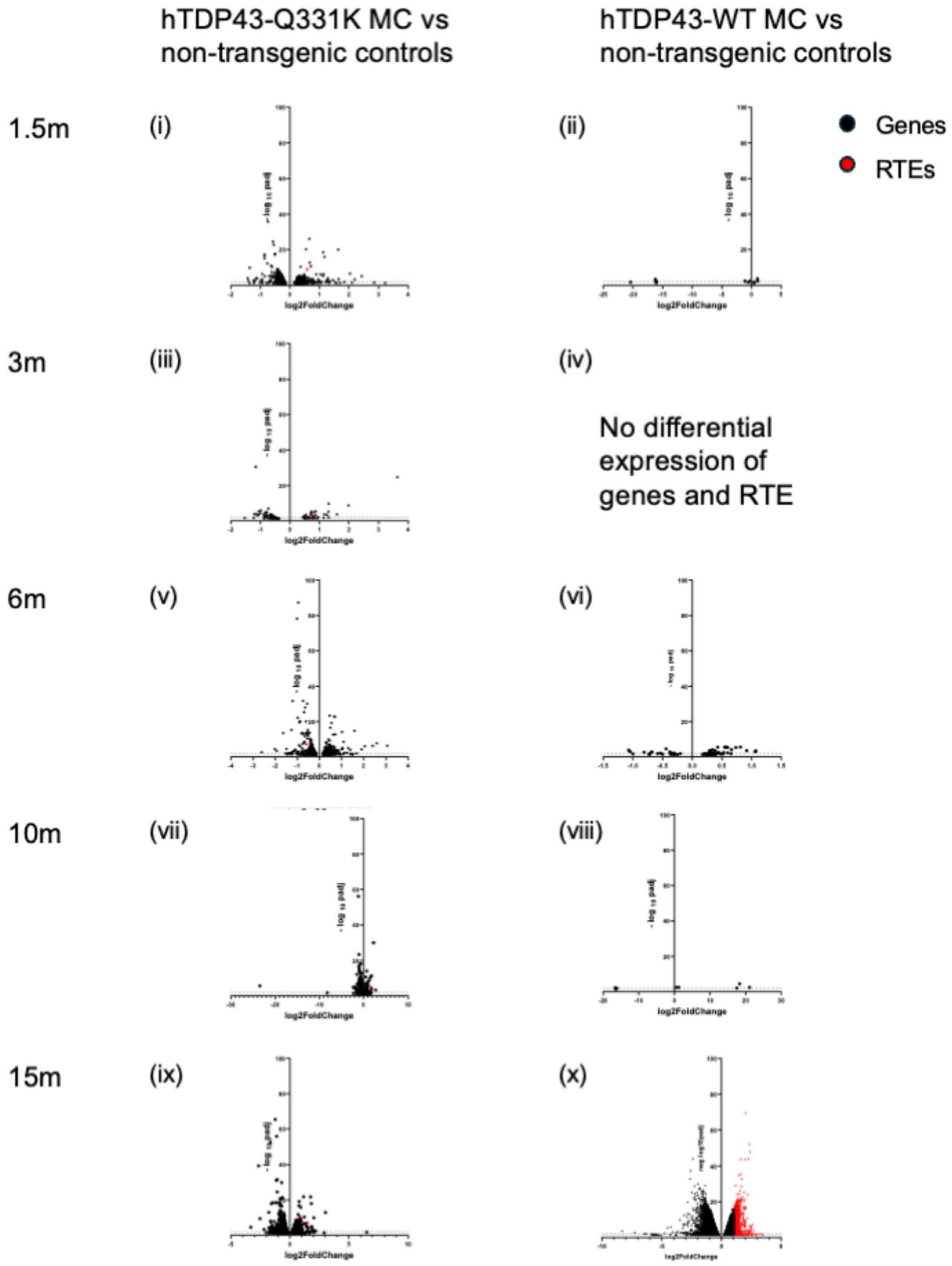

#### 3-month hTDP43-Q331K vs nTg: Genes upregulated

i)

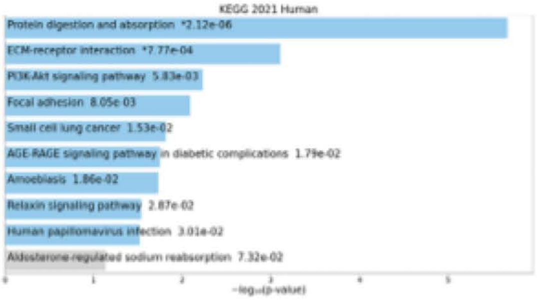


ii)

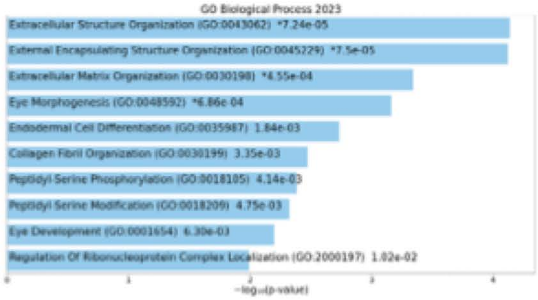


iii)

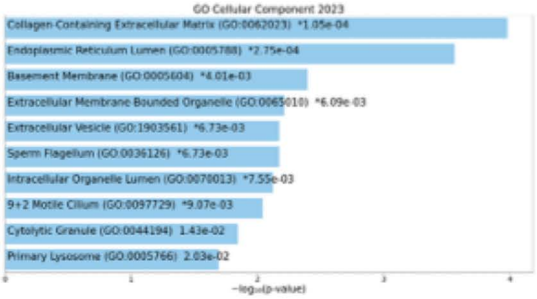


iv)

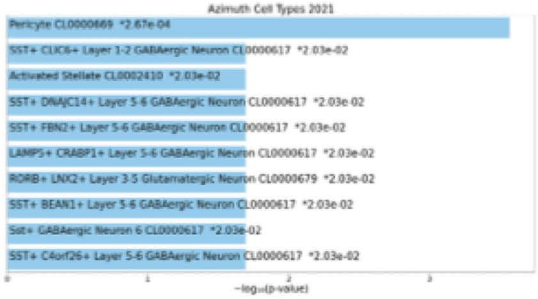


#### 3-month hTDP43-Q331K vs nTg: Genes downregulated

v)

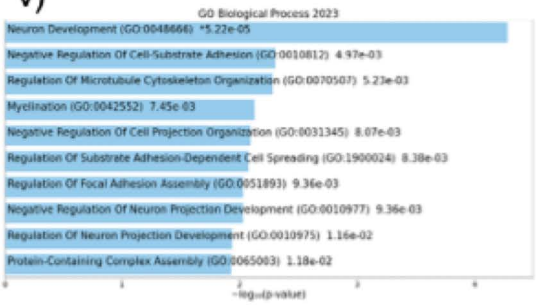


vi)

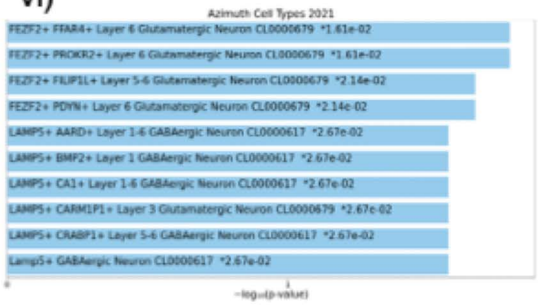

### 15month hTDP43-Q331K vs nTg: Genes upregulated

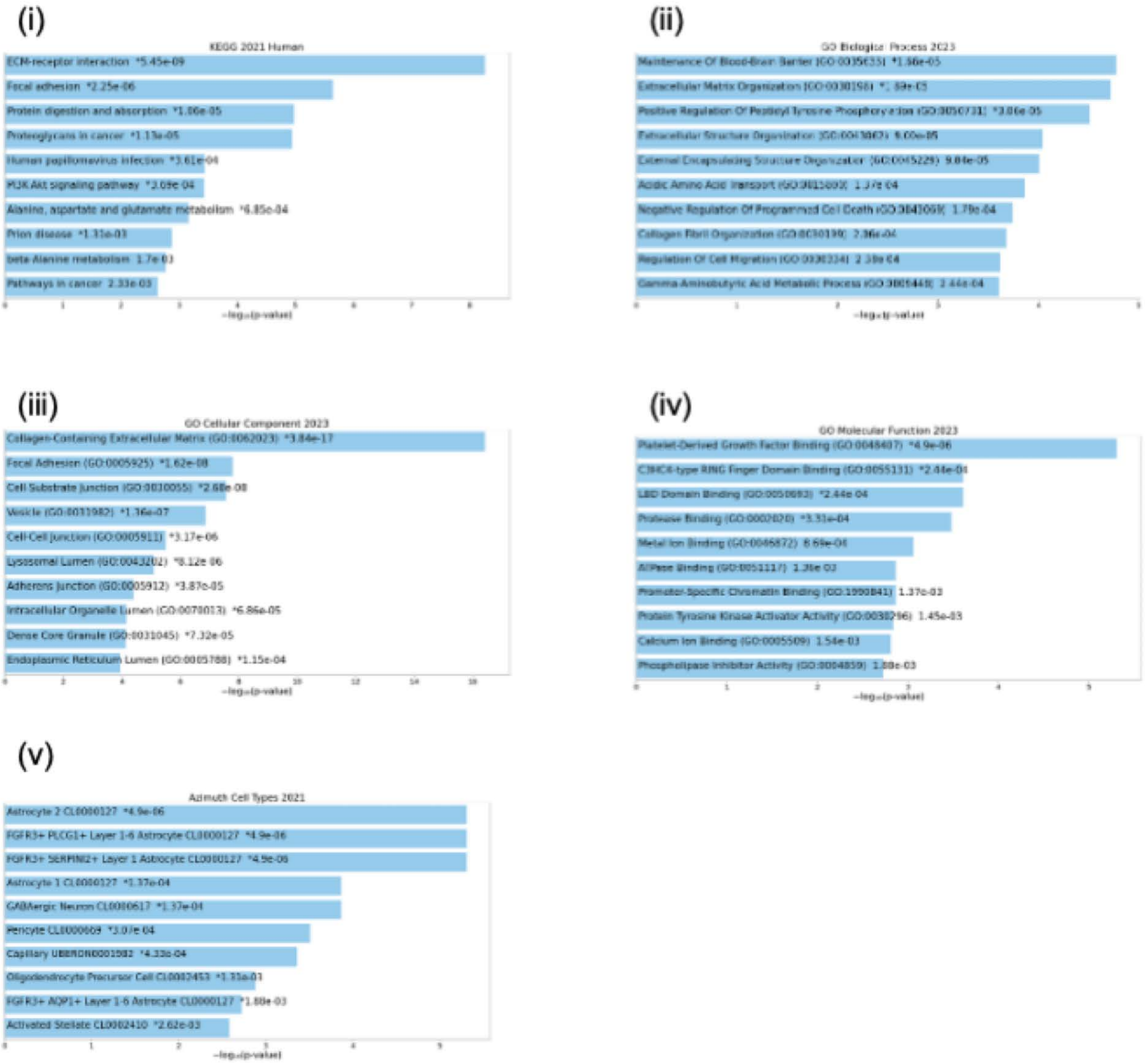

Supplementary Figure 2e

### 15month hTDP43-Q331K vs nTg: Genes downregulated

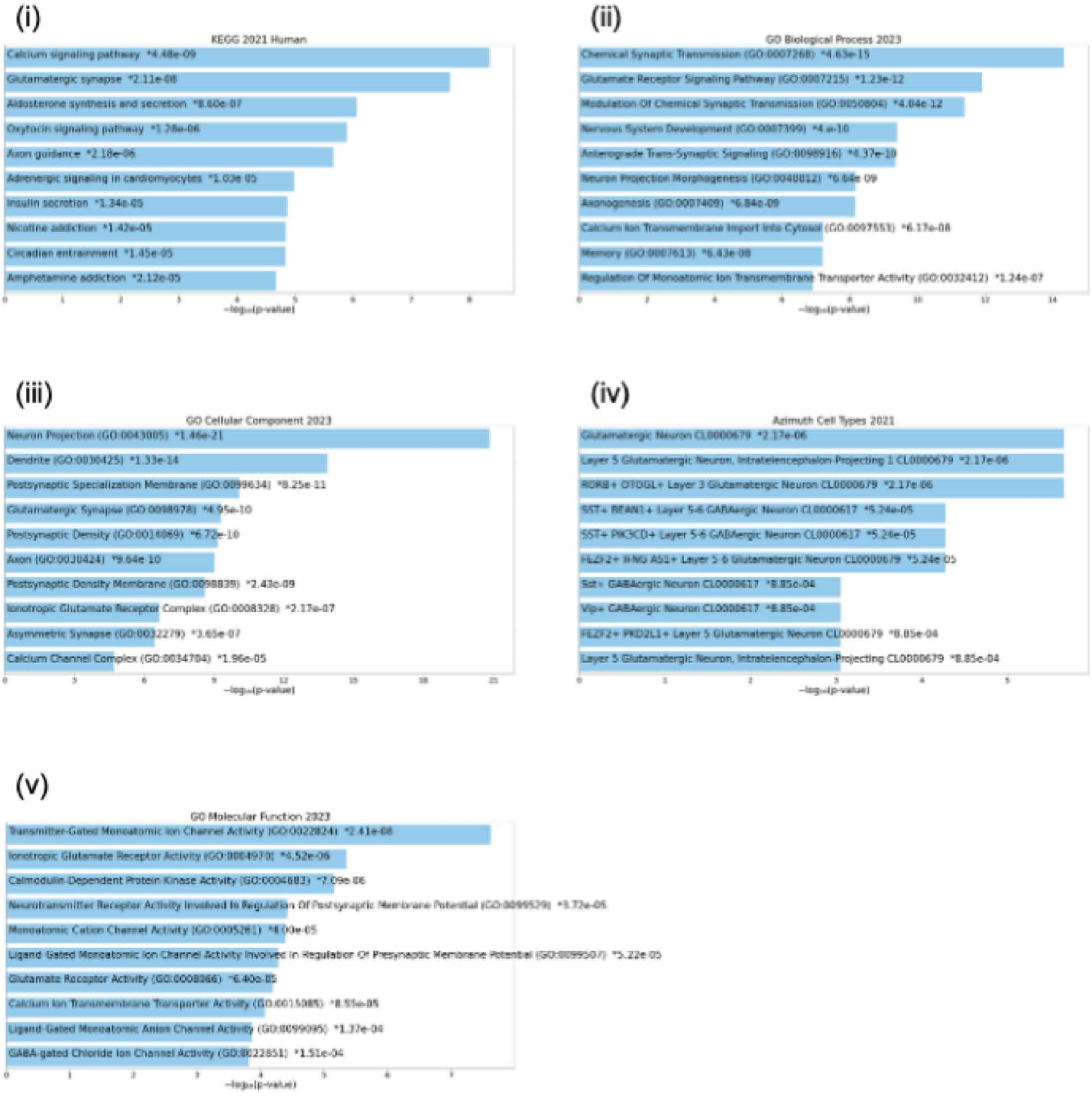

#### 15month hTDP43-WT vs nTg: Genes upregulated

(i)

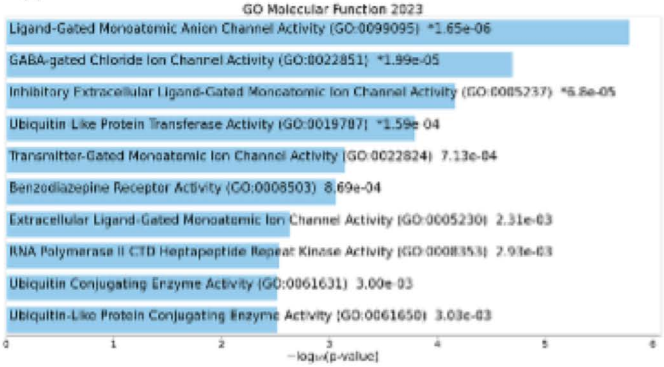


#### 15month hTDP43-WT vs nTg: Genes downregulated

(ii)

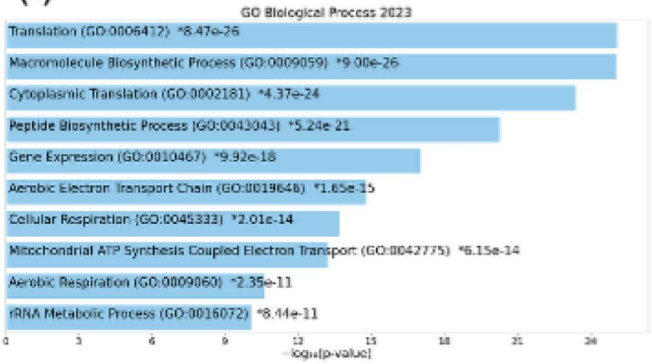


(iii)

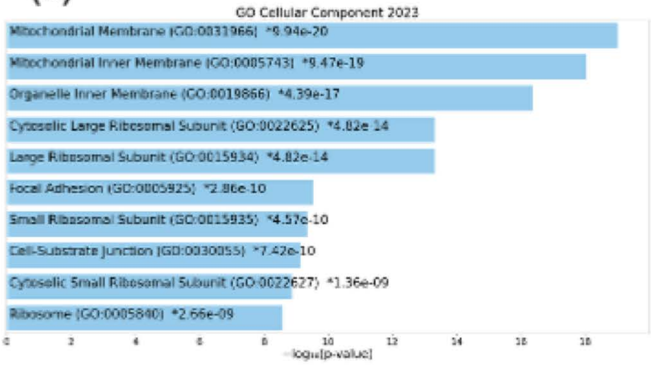


(iv)

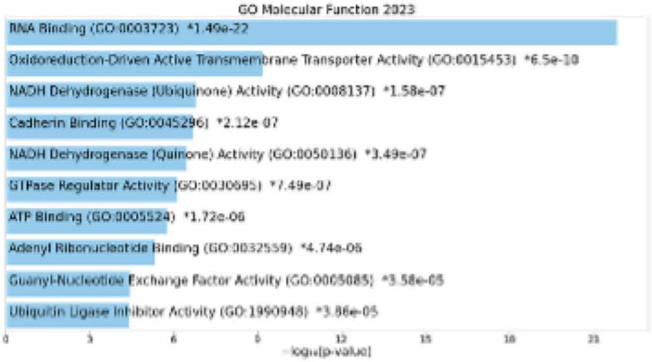


(v)

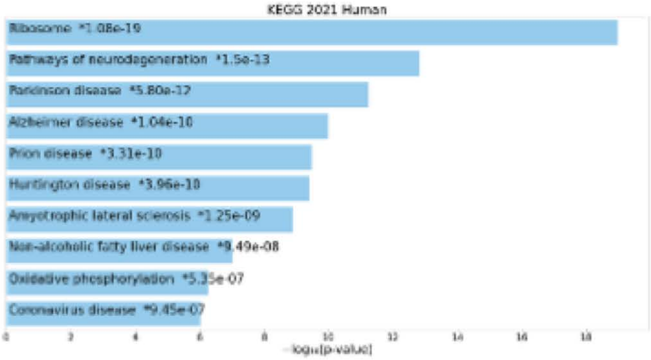

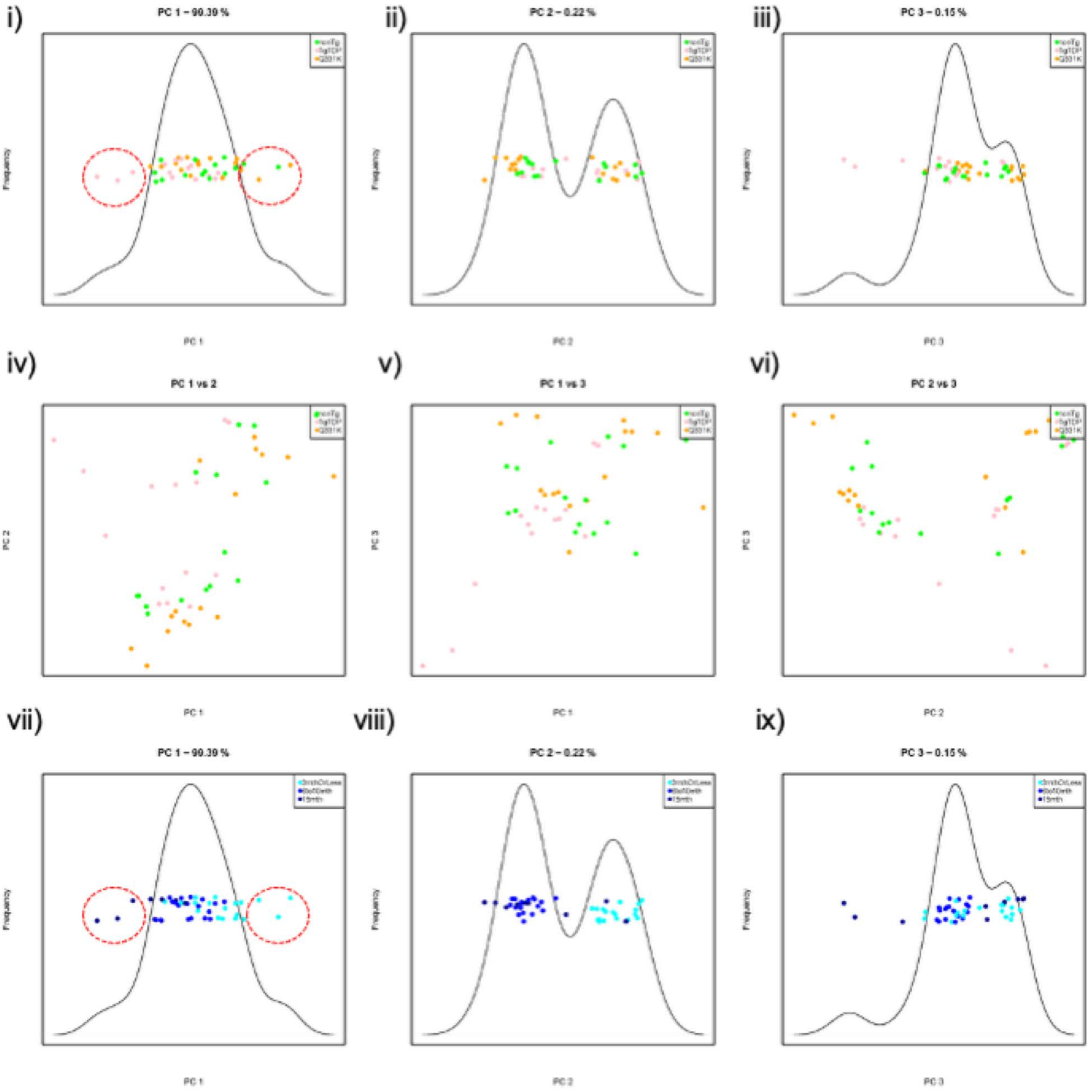
i)
PC 1 – 90.39 %
Frequency
PC 1
ii)
PC 2 – 0.22 %
Frequency
PC 2
iii)
PC 3 – 0.15 %
Frequency
PC 3
iv)
PC 1 vs 2
PC 2
PC 1
v)
PC 1 vs 3
PC 3
PC 1
vi)
PC 2 vs 3
PC 3
PC 2
vii)
PC 1 – 90.39 %
Frequency
PC 1
viii)
PC 2 – 0.22 %
Frequency
PC 2
ix)
PC 3 – 0.15 %
Frequency
PC 3

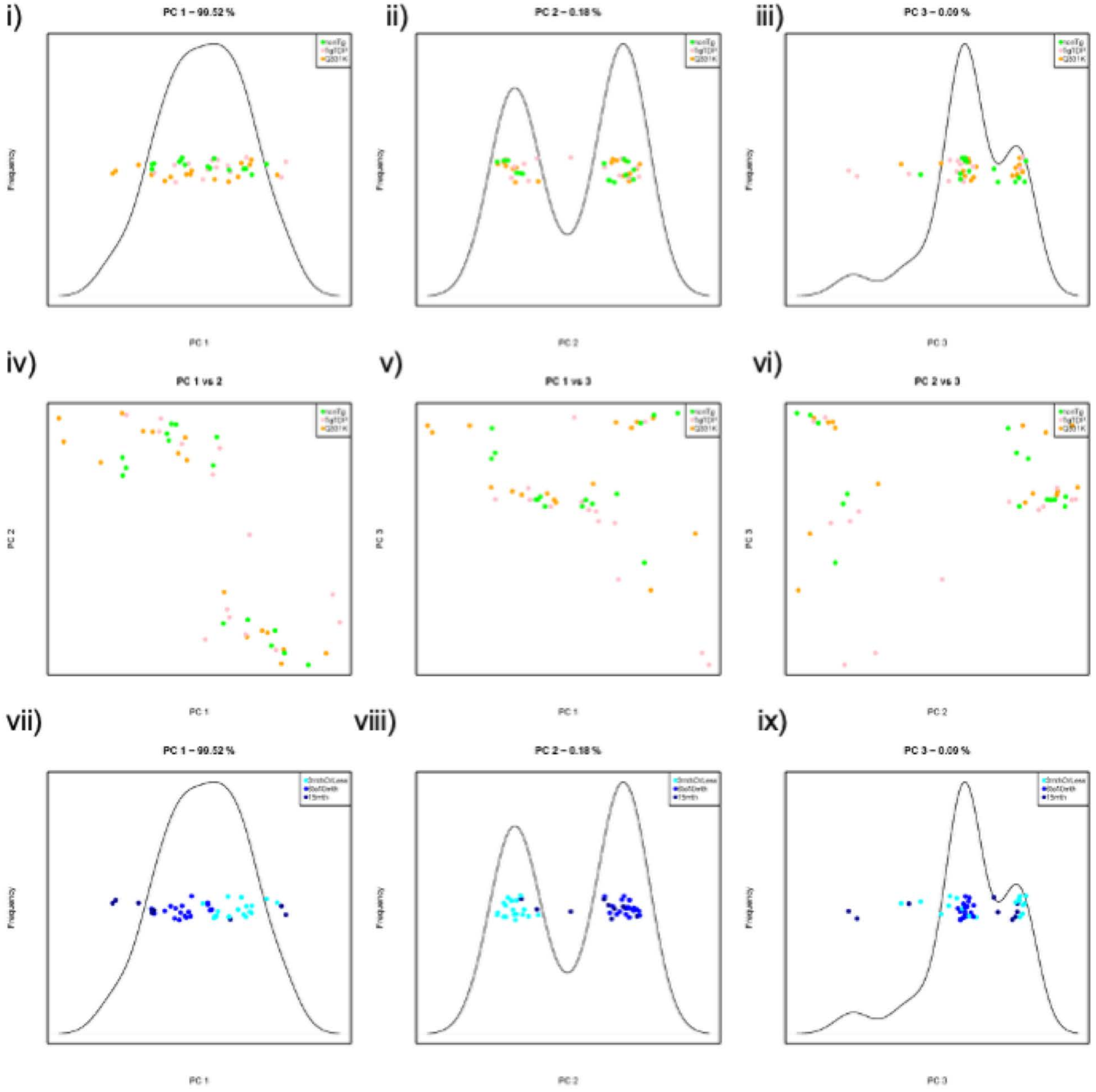
i)
PC 1 – 99.52 %
ii)
PC 2 – 0.18 %
iii)
PC 3 – 0.09 %
Frequency
PC 1
PC 2
PC 3
iv)
PC 1 vs 2
v)
PC 1 vs 3
vi)
PC 2 vs 3
vii)
PC 1 – 99.52 %
viii)
PC 2 – 0.18 %
ix)
PC 3 – 0.09 %

Supplementary Figure 2i

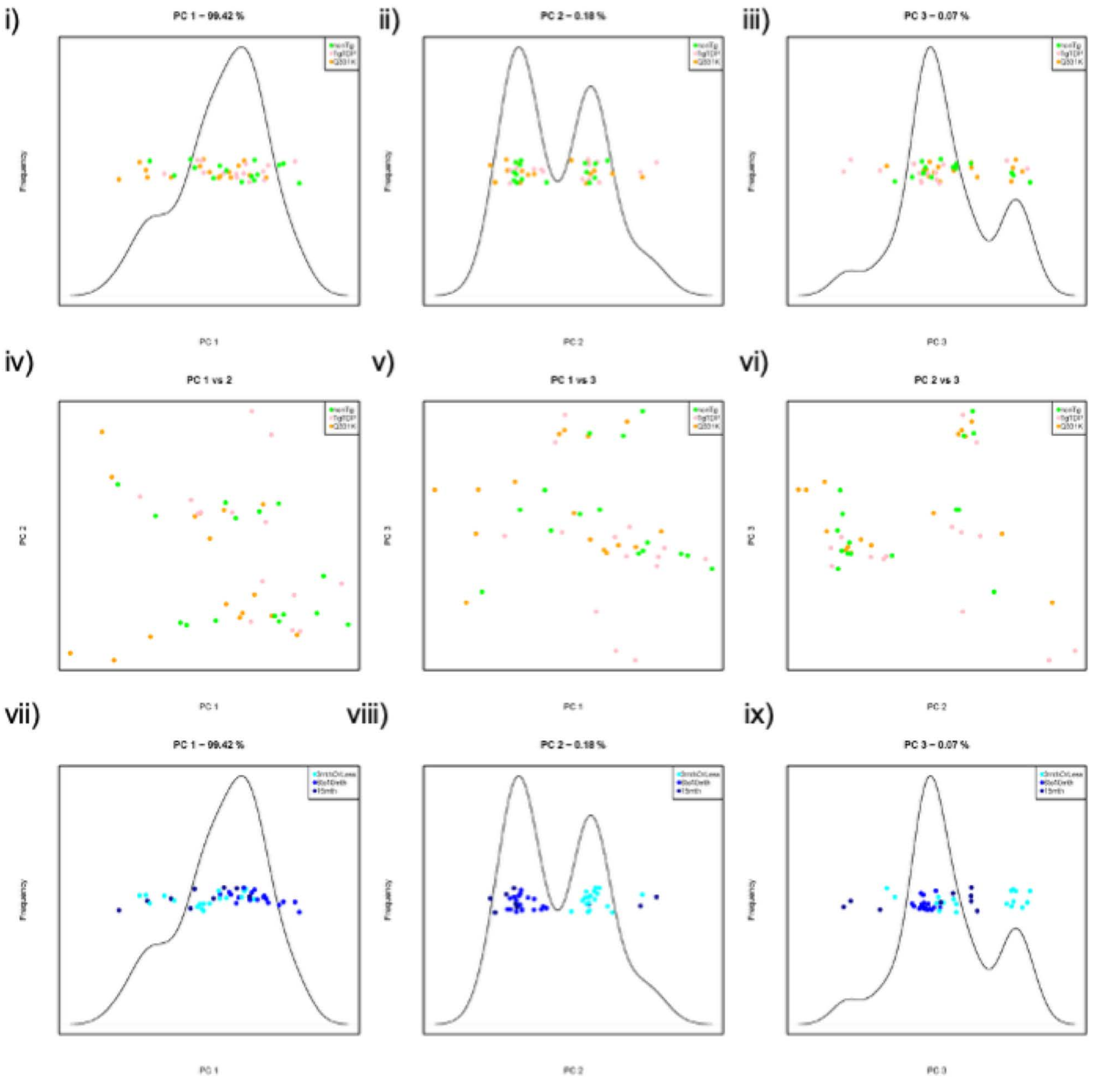

Supplementary Figure 2j

i) RTEs high during development

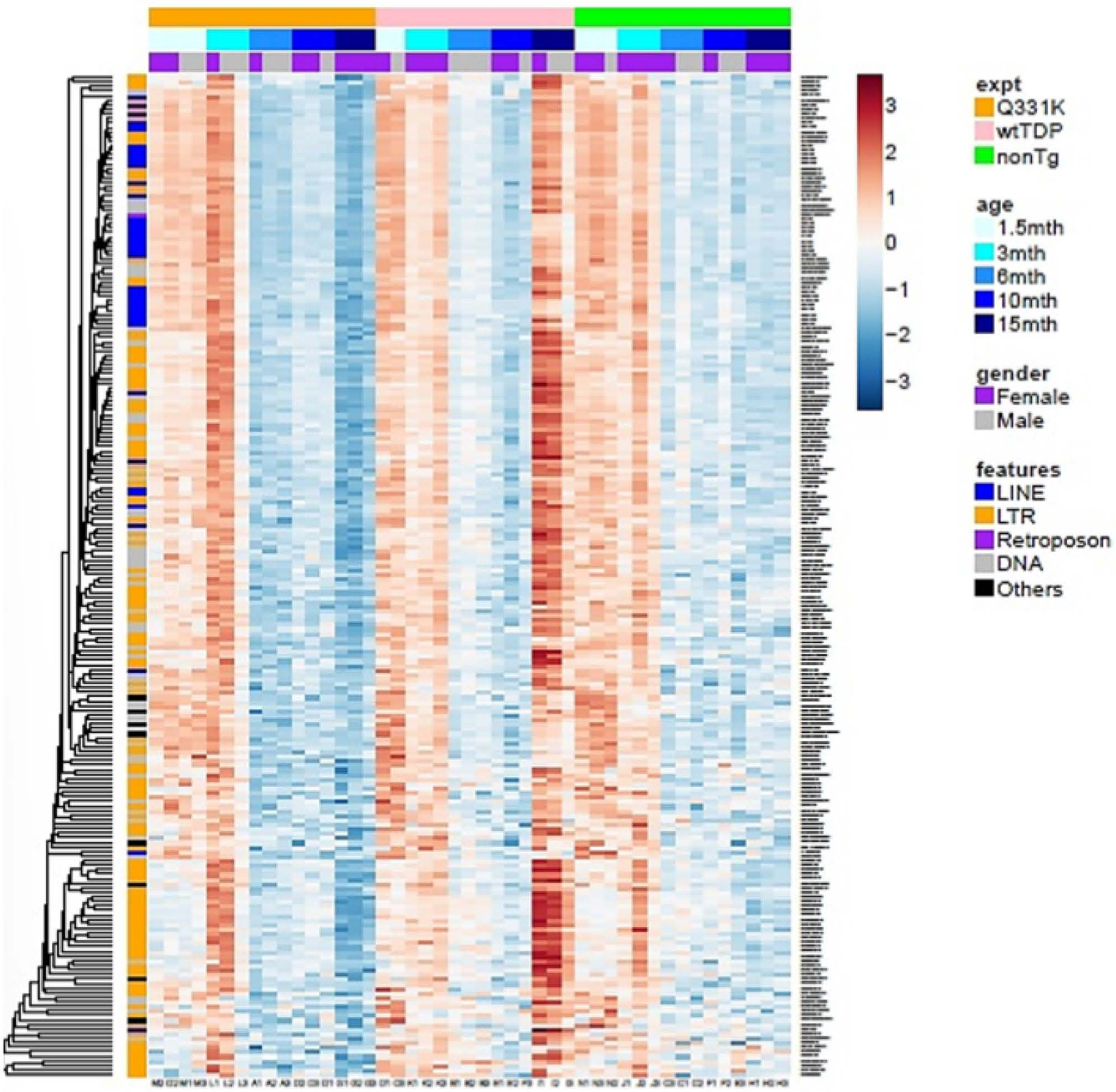

i) RTEs high in old animals

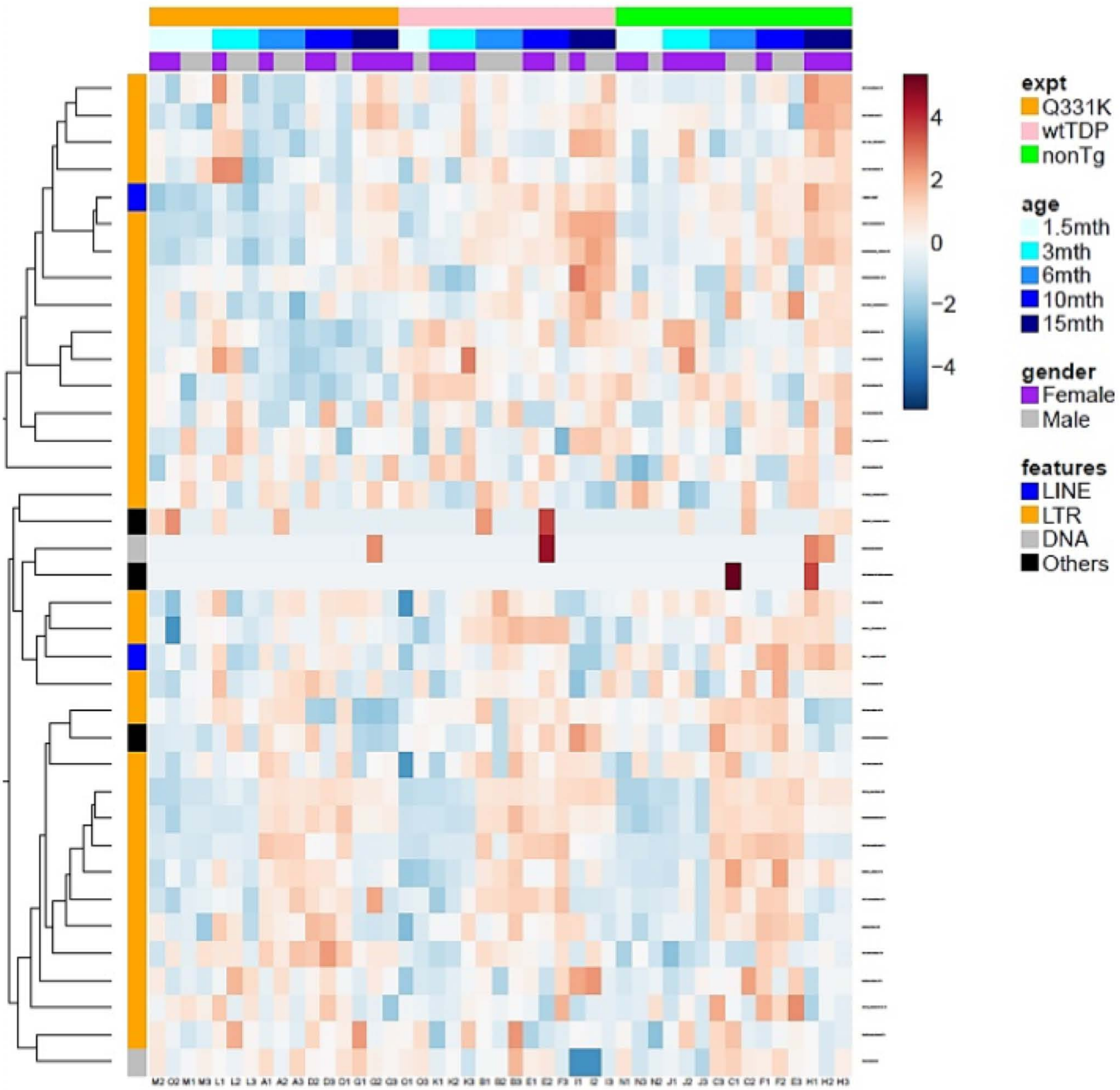

i) Significantly differentially expressed genes involved in the cell-cycle at 3 months in the hTDP43-Q331K motor cortex when compared to non-transgenic littermates

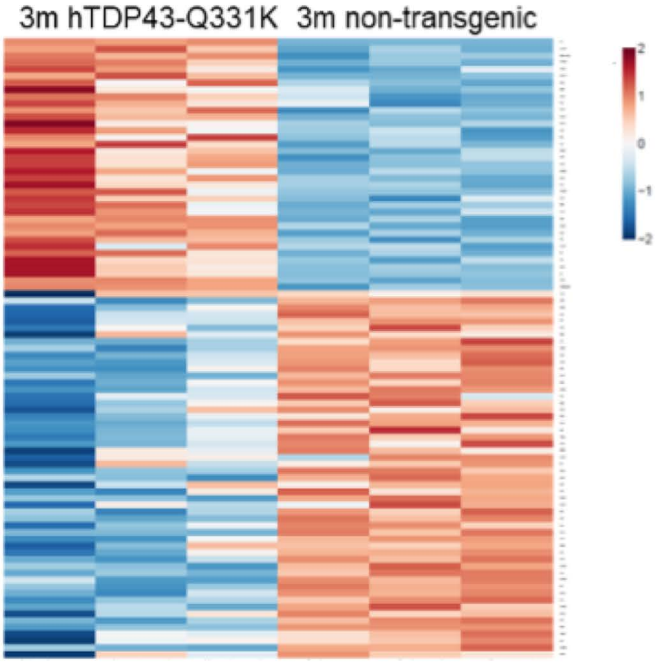


ii) Significantly differentially expressed genes involved in the cell-cycle at 15 months in the hTDP43-WT motor cortex when compared to non-transgenic littermates

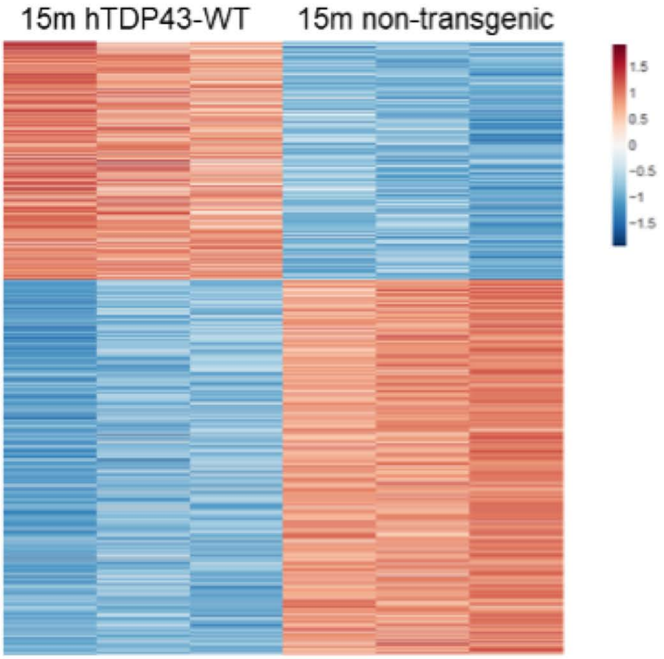

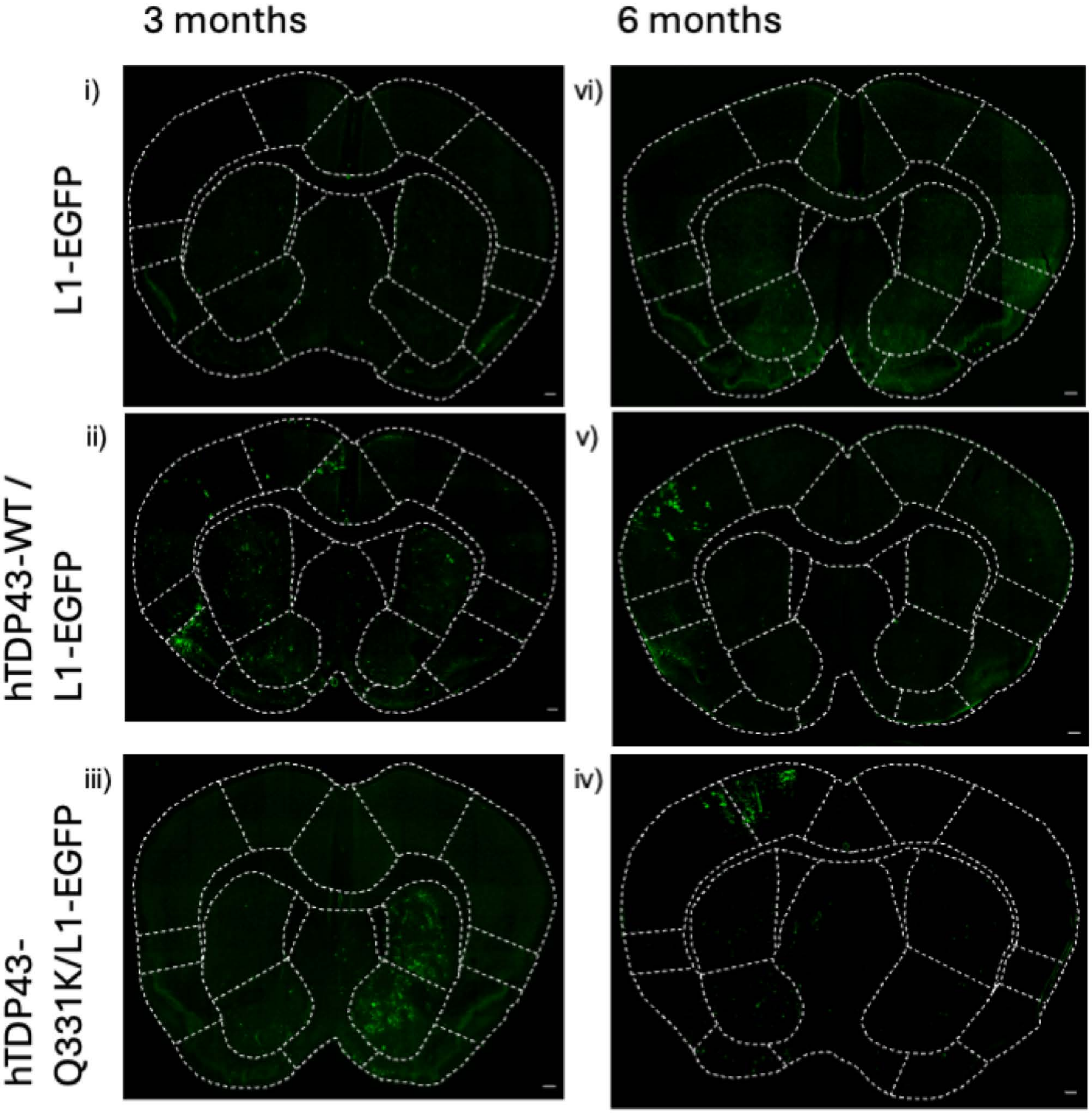
3 months
6 months
i)
vi)
L1-EGFP
ii)
v)
hTDP43-WT /
L1-EGFP
iii)
iv)
hTDP43-
Q331K/L1-EGFP

#### (i) 3-month-old motor cortex LINE1 replication clusters

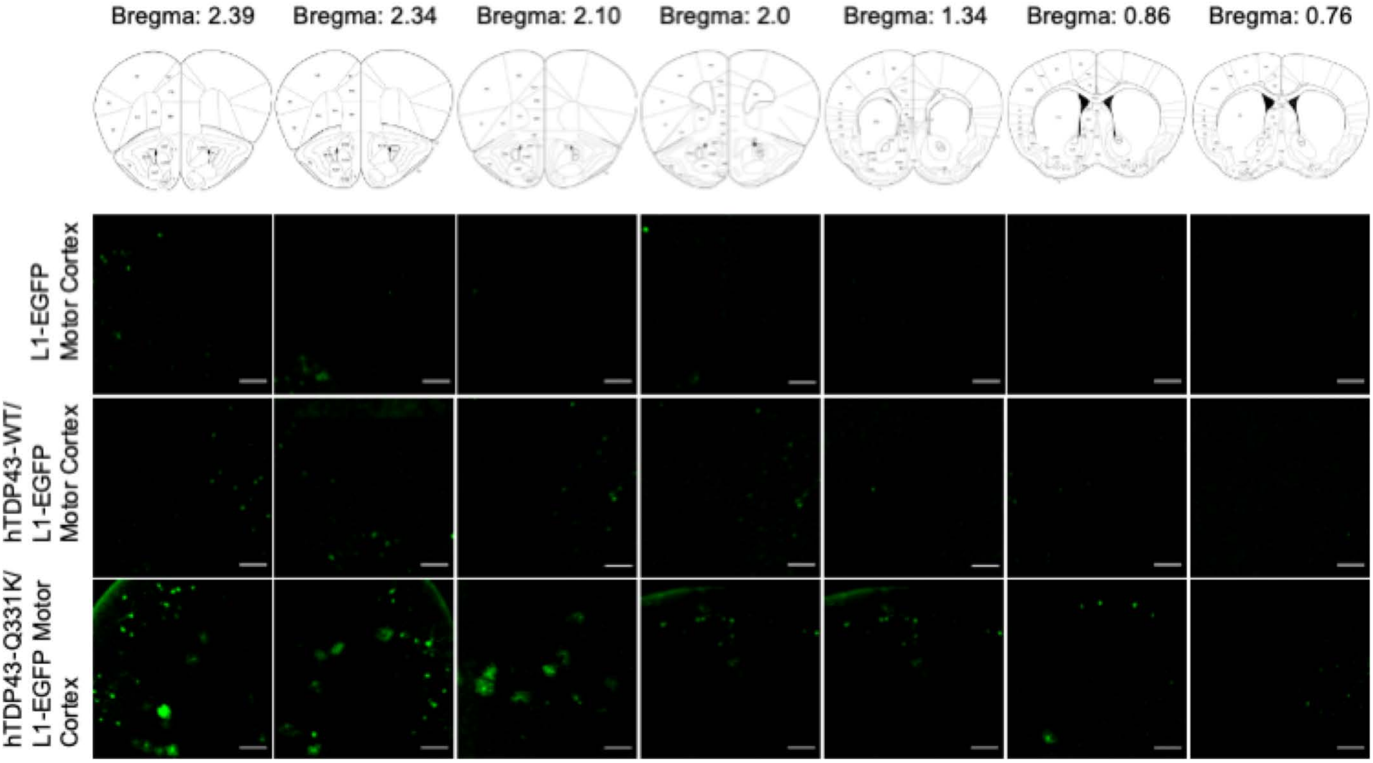


#### (ii) 6-month-old motor cortex LINE1 replication clusters

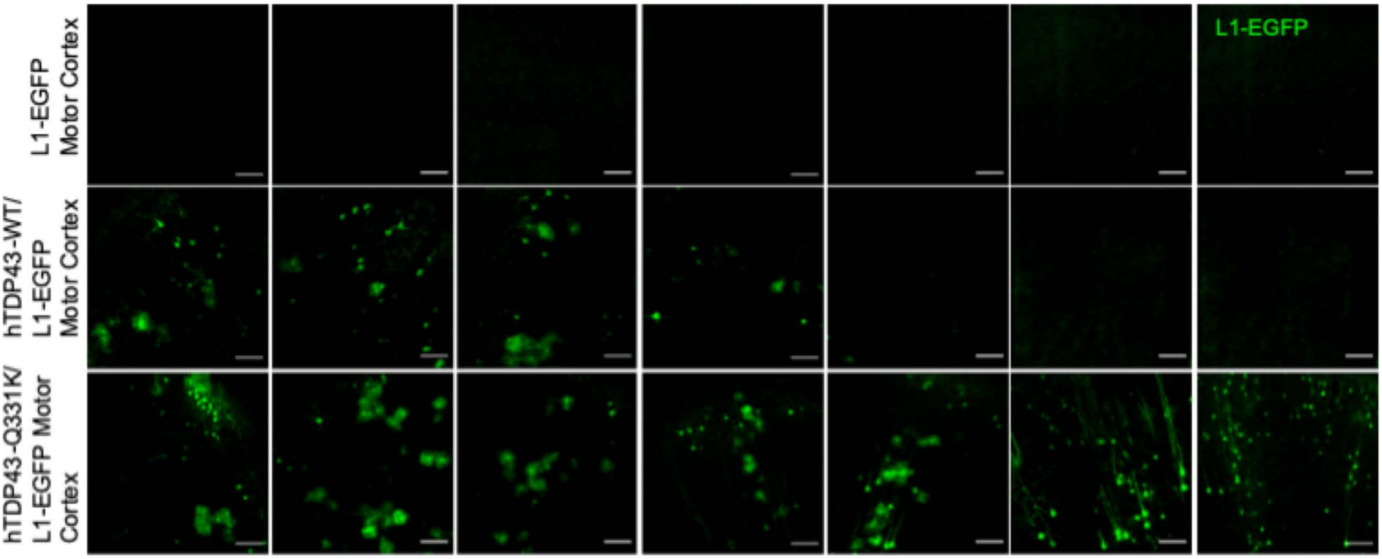

Supplementary Figure 4a

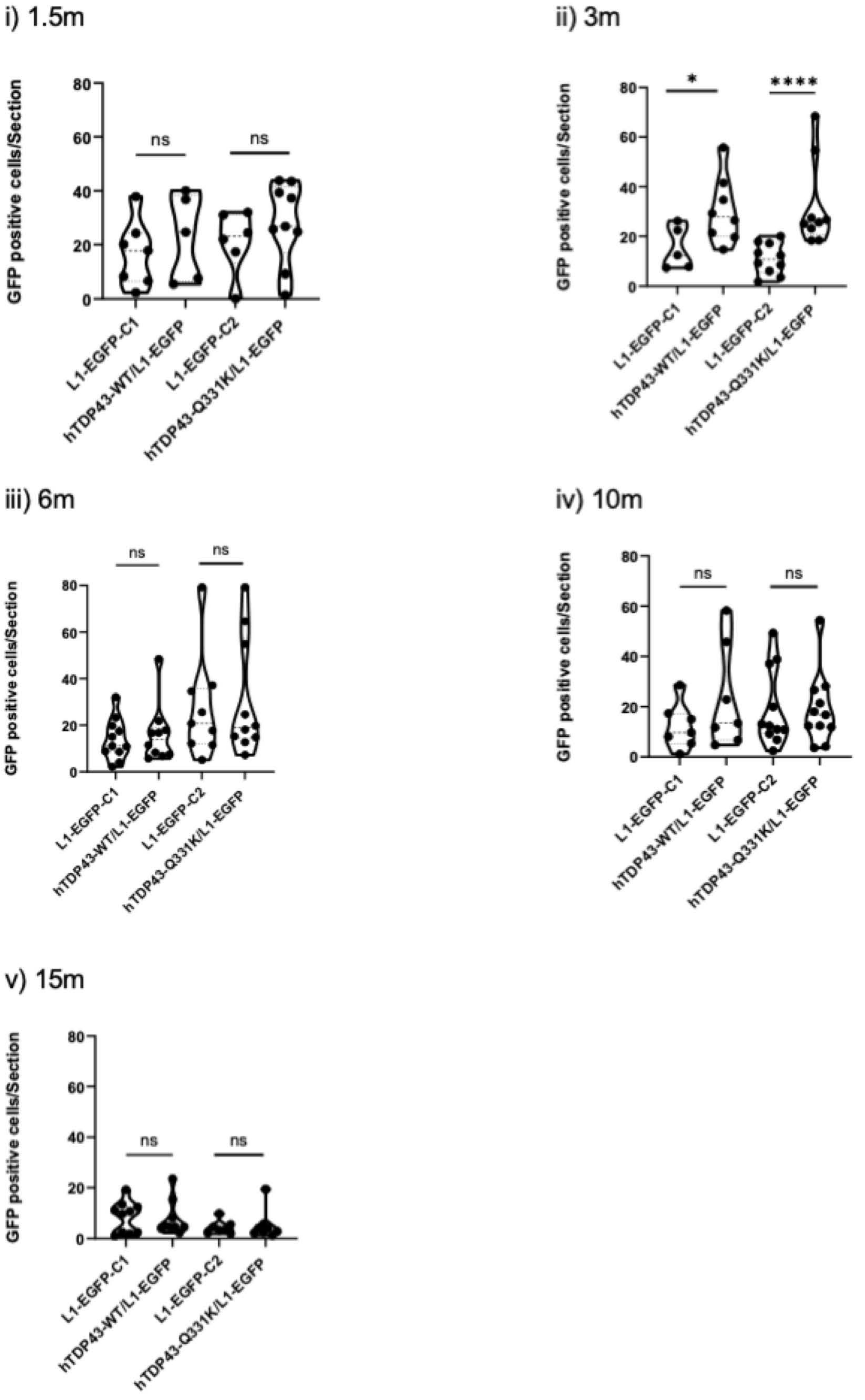

Supplementary Figure 4b

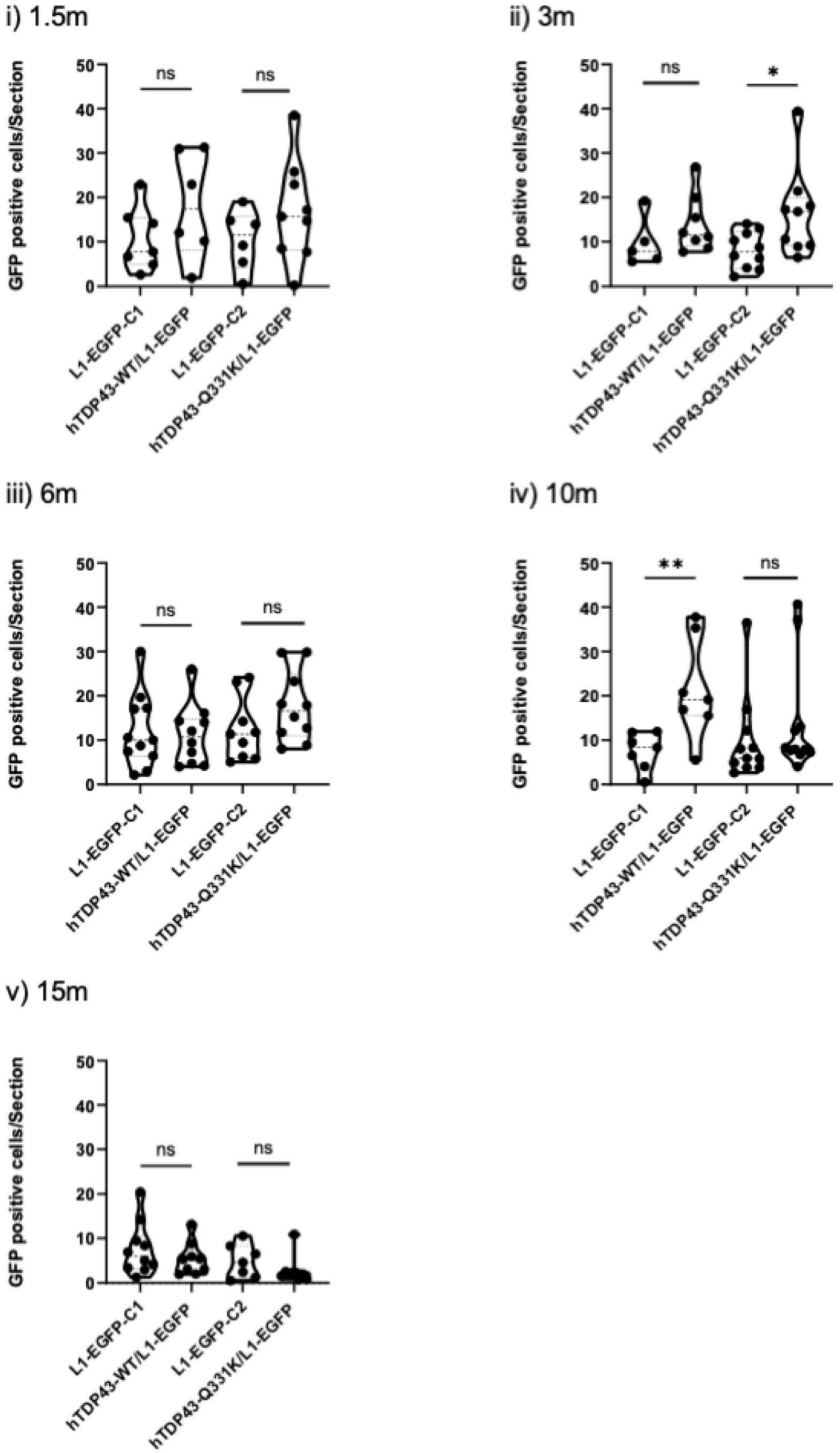

Supplementary Figure 4c

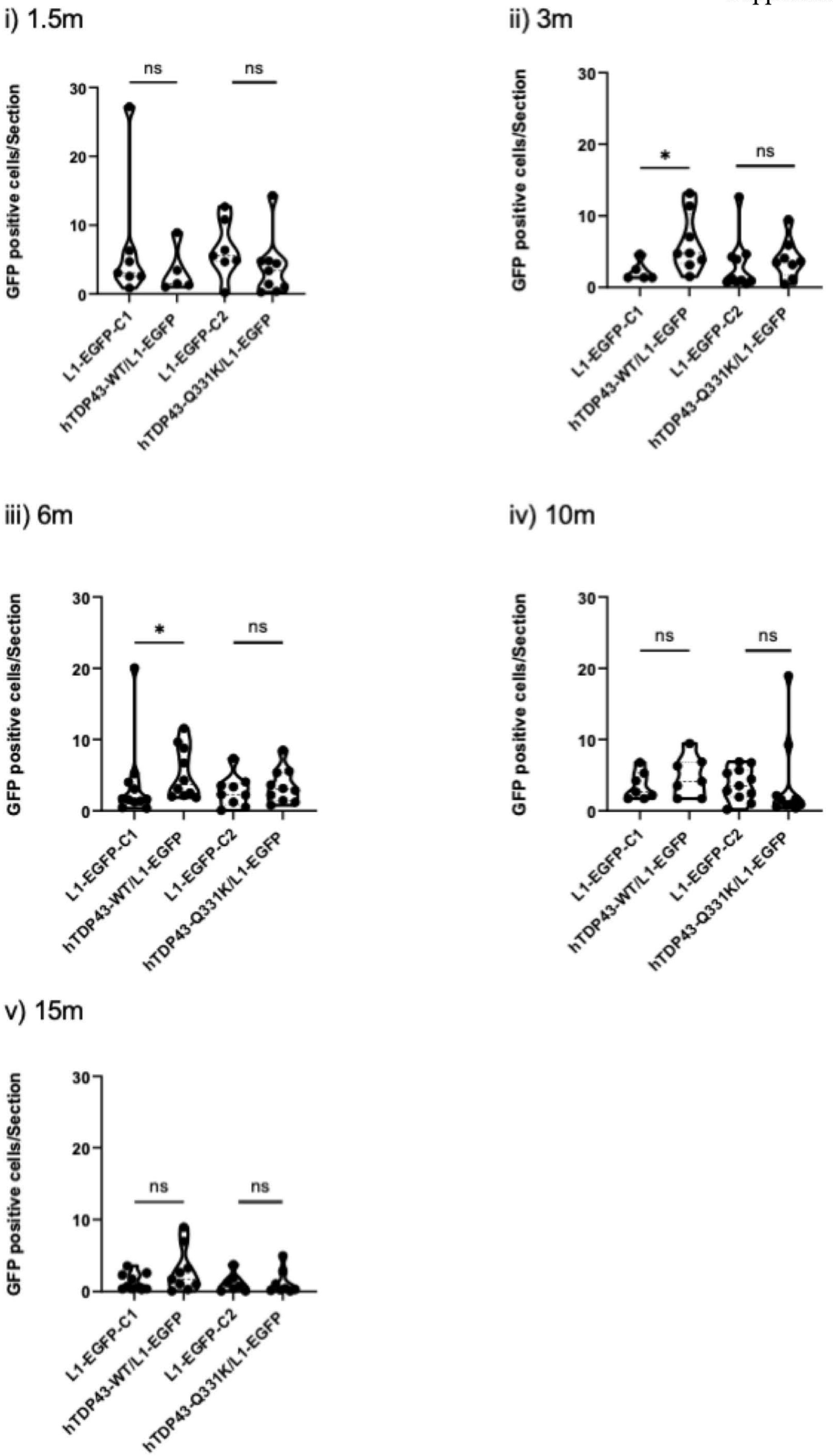
